# A toolkit to generate inducible and interconvertible *Drosophila* transgenes

**DOI:** 10.1101/2020.08.18.256461

**Authors:** Franz Wendler, Sangbin Park, Claire Hill, Alessia Galasso, Kathleen R. Chang, Iman Awan, Yulia Sudarikova, Mar Bustamante-Sequeiros, Sichen Liu, Ethan Sung, Gabrielle Aisabonoko, Seung K. Kim, Luis Alberto Baena-Lopez

**Affiliations:** Sir William Dunn School of Pathology. University of Oxford. South Parks Road. Oxfordshire, UK; Department of Developmental Biology, Stanford University School of Medicine, Stanford, CA, USA; Department of Medicine, and Stanford Diabetes Research Center, Stanford University School of Medicine, Stanford, CA, USA

**Author notes:** Authors with equal contribution (Franz Wendler, Sangbin Park).

**Keywords:** Gal4/UAS, QF/QUAST, LexA/LexAop, Flippase-FRT mediated recombination, Cre/LoxP-mediated recombination, Drosophila overexpression transgene

## Abstract

The existence of three independent binary systems for conditional gene expression (Gal4/UAS; LexA/LexAop; QF/QUAST) has greatly expanded versatile genetic analyses in the fruit fly *Drosophila melanogaster*; however, the experimental application of these tools is limited by the need to generate multiple collections of non-interchangeable transgenic fly strains for each inducible gene expression system. To address this practical limitation, we developed a modular vector that contains the regulatory elements from all three binary systems, enabling Gal4-, LexA- or QF-dependent expression of transgenes. Our methods also incorporate DNA elements that facilitate independent site-specific recombination and elimination of regulatory UAS, LexAop or QUAST modules with spatial and temporal control, thus offering unprecedented possibilities and logistical advantages for *in vivo* genetic modulation and efficient interconversion of overexpression transgenic fly lines.

## MAIN

The inducible Gal4/UAS gene expression system revolutionized genetic experimentation in fruit flies^1^. This binary genetic tool facilitates gene expression in larval and adult fly tissues with cellular specificity and temporal control^1^. The Gal4/UAS system relies on the production of the yeast Gal4 transcription factor from gene-specific regulatory enhancers and adjacent promoters^1^. The Gal4 protein can bind to cognate DNA upstream activating sequences (UAS), thereby inducing transcription of any DNA sequence of interest inserted downstream of the UAS^1^. The Gal4/UAS system has been instrumental in deciphering many biological processes, and has inspired the development of analogous conditional binary gene expression tools such as the LexA/LexAop and QF/QUAST systems^2,3^. The use of these bipartite gene expression tools, singly or in combinations, has greatly broadened conditional genetics in fruit flies^4^. Currently, each of these gene expression tools requires the generation of independent transgenic fly lines, limiting experimental interchangeability and combinatorial usage, as well as generating cost inefficiencies related to the production and maintenance of multiple transgene collections.

To circumvent these limitations, we have developed a versatile modular vector (MV, hereafter) that incorporates the gene-activating regulatory sequences for three binary gene expression systems (5x lexAop / 5x UAS / 5x QUAS) upstream of a multi-cloning site (**Fig. 1a**). This design was predicted to enable the *in vivo* expression of an inserted DNA sequence under the regulation of the specific transactivator chosen. To demonstrate the flexible function of the MV, we generated a transgenic fly line encoding for a synthetic protein containing the small peptide HA fused to a fragment of a split venus fluorescent protein (VC fragment)^5^. We subsequently intercrossed the new *MV-HA-VC* fly strain with fly lines individually expressing different transcriptional activators (*decapentaplegic-lexA*, *dpp-lexA*^6^; *spalt-Gal4*, *sal-Gal4*^7^; or *Hedgehog-QF*, *Hh-QF^8^*). Each of these intercrosses generated F1 larval progeny and restricted HA-VC expression within Gal4, LexA and QF-transcribing cells, respectively (**Fig.1b-d**). Furthermore, the use in parallel of two transcriptional drivers (*dpp-lexA/Hh-QF*; *dpp-lexA/sal-G4*) induced the concomitant expression of HA-VC within their cellular domains (**Fig. 1e,f**). Confirming the expression of our full-length HA-VC protein, immunostaining for HA and GFP largely colocalised in all of the cells expressing the MV construct (**Supplementary Figure 1b**). Similar results were obtained expressing a form of the initiator caspase Dronc tagged with V5 and VC peptides at the C-terminus (**Supplementary Fig. 1c-e**). In this case immunoprecipitation experiments followed by Western blotting confirmed the full-length expression of the Dronc-V5-VC (the expected size of Dronc-V5-VC is ~62KDa; **Fig. 1g**). These results demonstrated that the MV allows the expression of full-length cDNAs under the control of either separate or combined bipartite gene expression systems. We speculate that enhanced spatial and temporal precision will be achievable by incorporating Gal80 or QS, the respective repressors of Gal4/LexA or of QF^4^. These MV features can streamline genetic configurations for investigating gene function in cells with different origins, and for studying inter-organ cell communication.

**Fig. 1.**
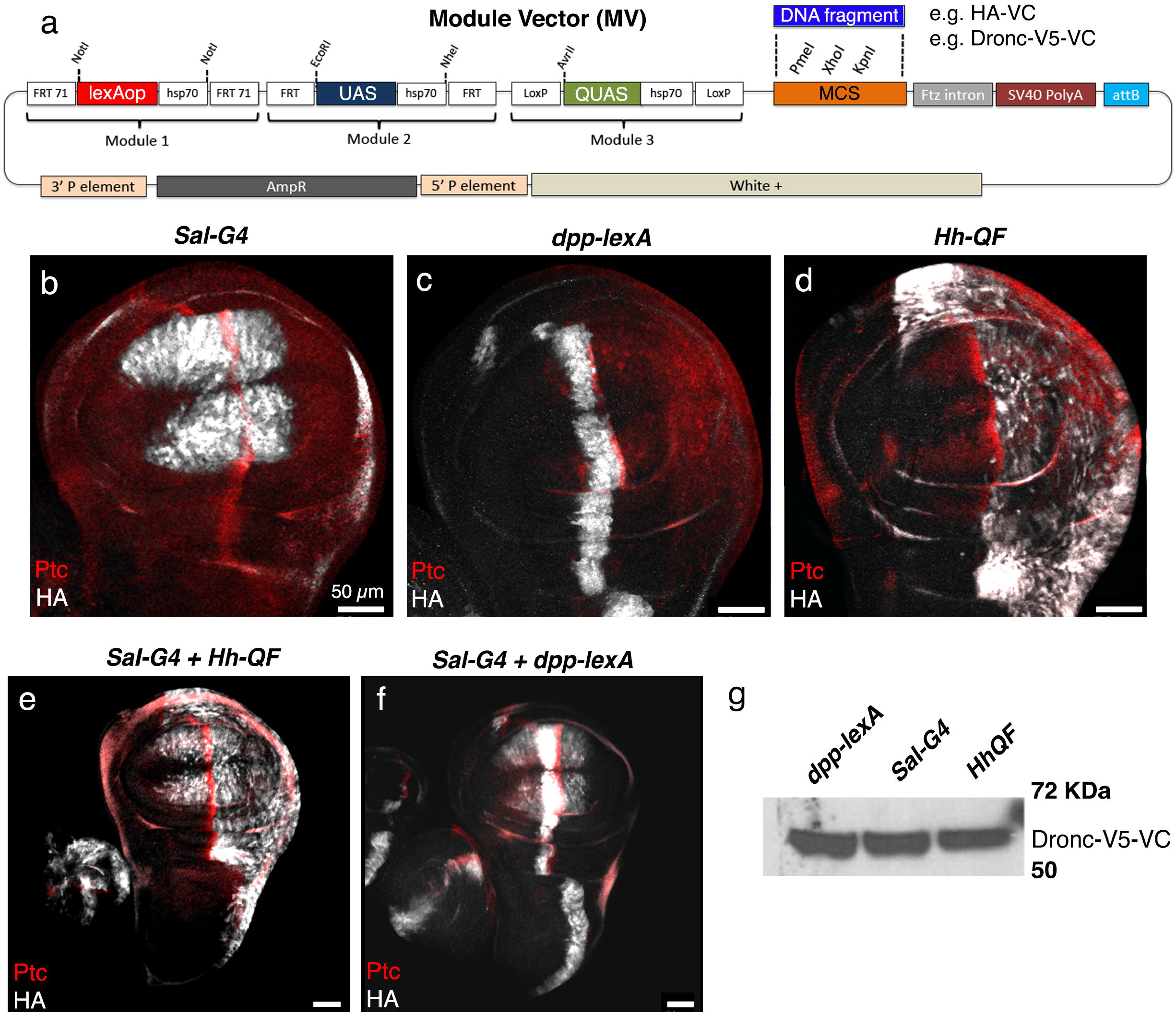
Features of the new Modular Vector (MV). **a)** Schematic showing the different features incorporated into the Modular-Vector (MV). **b-d)** Expression of the HA-VC (anti-HA, grey) within discrete cellular domains of the wing disc under the regulation of different transcriptional activators. **e-f)** Concomitant expression of the HA-VC in several cellular domains of the wing disc under the regulation of two different transcriptional activators. Patch expression (anti-Ptc, red) labels the confrontation between anterior and posterior cells of the wing disc. Scale bars represent 50 μm in the entire figure. **g)** Immunoprecitation followed by Western blotting of Dronc-V5-VC expressed under the regulation of several transcriptional activators; expected size of Dronc-V5-VC is around 62KDa.

Next, we explored the ability of our plasmid to generate loss-of-function phenotypes expressing short-hairpin RNA interference constructs. To this end we cloned short inverted repeats for targeting the gene *wingless* (flybase reference number = HMS00844) in the multicloning site of MV. Importantly, a TRIP line expressing this construct (HMS00844) was previously shown to downregulate the expression of Wg *in vivo* under the regulation of patch-Gal4^*9*^. However, our construct failed to induce noticeable *wg* downregulation or *wg* mutant phenotypes upon overexpression with several drivers (e.g. *en-Gal4*, *Hh-QF* and *dpp-lexA*), Similar unexpected results were obtained using UAS-Dicer to enhance the efficiency of the knockdown in combination with *en-G4* and *nubbin-Gal4*, respectively. Although these preliminary experiments suggest that further optimization of our plasmid is likely needed to support robust RNA interference (e.g. expansion of the number of activation sequences), MV is still an excellent template for a wide range of biological applications requiring gene overexpression.

*Drosophila* genetics has benefited from three independent mechanisms of sequence specific genomic DNA recombination, especially the Flippase/FRT (Flp/FRT) system^10^. This system capitalises on Flp DNA recombinase activity that targets Flp-recognition sites (FRT sequences)^10^. Recently, directed mutagenesis of the ‘wild type’ Flp enzyme (hereafter, Flp^WT^) and the FRT-recognition sites generated the mFlp^5^/mFRT 7.1 system^11^. Another alternative, the Cre/LoxP system, relies on Cre-mediated recombination of LoxP-recognition motifs^12^. Because no cross-reactivity has been reported between these three recombination systems^11,13^, gene regulatory elements in the MV were flanked by specific pairs of these recombination sequences (**Fig. 2a**) to investigate if sequence-specific excision of gene-regulatory sequences could be achieved. Specifically, the UAS sites were flanked by FRT (Flp sensitive), the QUAST repeats by loxP (Cre sensitive) and the LexAop site by mFRT 7.1 (mFlp^5^ sensitive: **Fig.2a**). Upon expression of the cognate recombinase (Flp^WT^, Cre or mFlp^5^), the specific regulatory components were successfully excised from the genome in somatic cells (**Fig. 2a-c**; Methods). This facilitates the termination of the overexpression (GFP negative clonal cell populations indicated by red arrowheads in the right panels of **Fig. 2a-c**) induced by each transcriptional activator with temporal precision and the generation of genetic mosaics within somatic tissues.

**Fig. 2.**
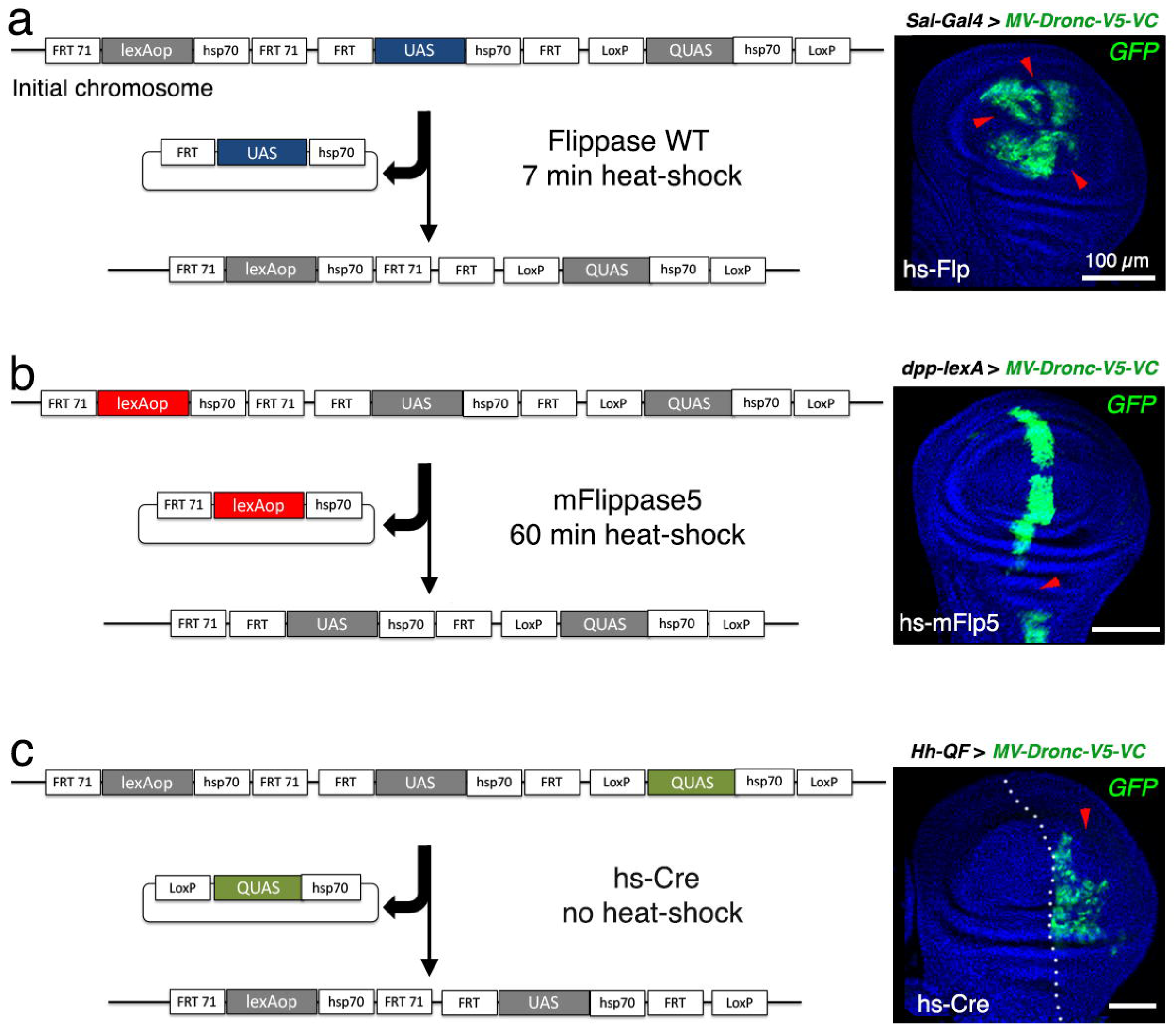
Generation of genetic mosaics in somatic tissues upon genomic elimination of specific gene regulatory elements included in the MV plasmid. **a-c)** Clonal elimination from the genome of 5X UAS (a), 5X LexAop (b), and 5X QUAS (c) binding sites upon random exposure to wild type Flippase (a), mFlippase5 (b) and Cre (c), respectively; anti-GFP is used to detect the expression of Dronc-V5-VC (green) and Dapi (blue) labels the DNA; red arrowheads indicate the lack of expression of Dronc-V5-VC in the expression domain of the different drivers (Gal4, LexA and QF). Compare the expression of Dronc-V5-VC (green) within this figure to Supplementary Fig. 1. Notice that the hs-Cre transgene is basally expressed in random cells without heat-shock induction^16^. Scale bars represent 100 μm in the entire figure.

Next, we assessed whether gene *cis*-regulatory elements could be excised in the germline of a founder MV fly strain; if so, this would provide a means to efficiently generate derivative strains with permanently altered gene induction capacities. We intercrossed the MV-*Dronc-V5-VC* founder line with flies expressing Flp^WT^ from a *Beta2-tubulin* promoter (*BTub2*, hereafter) (**Fig. 3a**) to direct highly efficient FRT-dependent recombination in the male germline^14,15^. F1 male progeny containing both transgenes (M*V-Dronc-V5-VC* and *Btub2-Flp^WT^*) were subsequently intercrossed with females harbouring both the *sal-Gal4* and *dpp-lexA* driver transgenes (**Fig. 3b**). Remarkably, 95% (40/42 wing discs from 26 larvae) of the F1 (5 independent males were used to perform the experiment) generated larvae that did not express the Dronc-V5-VC construct in the *sal-Gal4* domain but retained the activation in the *dpp-lexA* territory (**Fig. 3c**). These results demonstrated that most of the males expressing Flp^WT^ in the germline permanently excised the UAS regulatory repeats without compromising the LexA-binding sites. Comparable results were obtained when Cre was expressed under the regulation of the *Btub2* promoter (**Fig. 3b, 3c**).

**Fig. 3.**
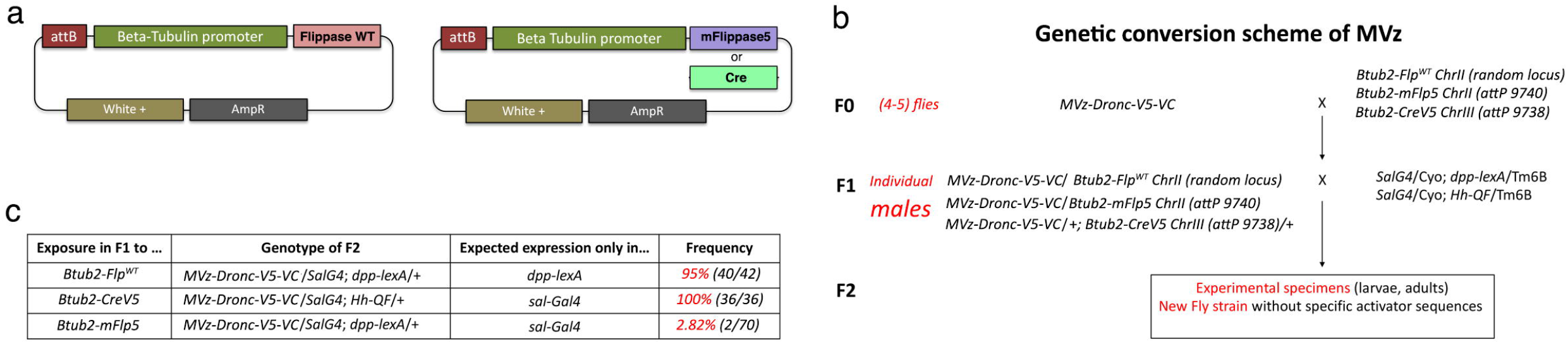
Genetic strategies to eliminate specific regulatory elements in the germline to generate new fly lines with selected gene activation sequences. **a)** Schematic showing the plasmids specifically designed to express different recombinases under the regulation of *Btub2* promoter. **b)** Genetic scheme of intercrosses to permanently eliminate regulatory components of MV constructs in the male germline; this feature can be used to create new transgenic flies containing only a subset of activating binding sites. **c)** Frequency of germline excision of each gene regulatory element upon exposure to different recombinases.

Unlike the results obtained when using Flp^WT^ or Cre, we found that intercrosses using flies expressing *Btub2-mFlp^5^* only affected expression of the Dronc-V5-VC in approximately 2.82% of analysed larvae (2/70 wing discs from 35 larvae). These results are likely due to a low recombination frequency mediated by mFlp^5^ in the male germline^15^, which could be alleviated by creating in the future new fly strains with enhanced Flp5 expression within male gametes (e.g. nanos-Gal4 UAS-Flp5). Despite this potential limitation, our findings confirm the suitability of the MV to efficiently obtain new fly strains starting from a single founder MV-line through a simple intercross strategy for at least two of the gene regulatory elements included in the MV plasmid (**Fig. 3b**). Moreover, the efficient germline excision of specific regulatory elements allows use of novel MV fly strains in combination with previously created lines in public repositories for multiplexed.

In summary, we have generated a next-generation modular vector that offers unprecedented experimental opportunities, whilst providing enhanced versatility to conduct targeted genetics in *Drosophila*. This expansion of the fruit fly genetic toolkit allows development of interconvertible fly collections of overexpression transgenes with a substantial reduction of logistical costs, including experimenter time and effort. Resulting lines retain specific gene regulatory elements of interest and can be rapidly interconverted through a simple scheme of intercrosses to allow altered inducible transgene regulation.

## MATERIAL AND METHODS

### Molecular cloning

All PCRs were performed with Q5 High-Fidelity polymerase from New England Biolabs (NEB, M0492L). Standard subcloning protocols were used to generate all the DNA plasmids (see details below). Genomic DNA was extracted via standard protocols was used as a template to amplify different DNA fragments. Transgenic flies harbouring the new transgenes were obtained by attP/attB PhiC31-mediated integration. Some of the transgenes were generated by Bestgene Inc, whilst others were generated in house by Sangbin Park. Fly strains generated will be deposited at the Bloomington Stock Centre. Whilst resources are transferred to Bloomington, reagents will be provided upon request.

#### Generation of the Module Vector (MV)

The different modules included in the vector were first designed *in silico* using the SnapGene software. Each module contains 5 repeats of each upstream regulatory sequence type, followed by an hsp70 minimal promoter. Each module was also flanked by specific recombination sites (see diagram Figure 1). This fragment was synthesized as a large DNA fragment by GENEWIZZ and subcloned in the PUC57 vector. The modular construct was then extracted from PUC57 as a SpeI-KpnI fragment and subcloned into a UASt-attB-w+ vector previously digested with NheI-KpnI. PmeI, XhoI, KpnI are potential unique restriction sites available for conventional cloning downstream of the gene regulatory elements (see diagram Figure 1). Sequence of the plasmid will be provided upon request until the vector is deposited in a public repository.

#### Generation of Beta2-Tubulin Flippase 5 and Cre transgenic flies

We extracted genomic DNA and amplified a Beta2-tubulin promoter via PCR using the following primers:

Forward primer Beta2-tubulin promoter:

*5’ ttattatccctaggcagctgtggactcctcattgtagg 3*’

*Reverse primer* Beta2-tubulin promoter:

*5’ aaatttaatctgcaggcggccgcgaattcaagcttcgcccctttttcacaccg 3*’

Convenient restriction sites for cloning were placed at the 5’ (AvrII) and 3’ (NotI and EcoRI) end of the PCR product. We then digested a UAS-attB-w+ vector with NheI and EcoRI, thus replacing the UAS repeats with the Beta2-tubulin promoter but keeping the rest of the plasmid backbone. The PCR was digested with AvrII and EcoRI and ligated with the aforementioned backbone, creating an intermediate plasmid (Beta2-tubulin-attB plasmid) suitable to subclone the cDNA of the Flippase-5 and Cre. The Flippase5 cDNA was amplified via PCR from a plasmid vector kindly provided by Iris Salecker using the following primers:

Forward primer Flippase-5:

*5’ attacagttGCGGCCGCatgccacaatttgatatattatgtaaaacacc 3*’

*Reverse primer* Flippase-5:

*5’ AAtATAaaggcctTctagattatatgcgtctatttatgtagg 3*’

The NotI and StuI restriction sites were conveniently placed in the PCR product of the Flippase-5 to facilitate the cloning in the Beta2-tubulin-attB plasmid. Before ligation, the Beta2-tubulin-attB plasmid and the Flippase-5 PCR product were digested with NotI and StuI.

To generate the Beta2-tubulin-CreV5-attB-w+ vector, we replaced the UAS repeats included in a UAS-Crev5-attB-w+ plasmids previously generated in the laboratory by the Beta2-Tubulin minimal promoter. To that end and prior ligation, we digested the UAS-CreV5-attB-w+ plasmid and the Beta2-Tubulin minimal promoter PCR product with NheI-EcoRI. Notice that the Cre cDNA was tagged with the V5 epitope.

#### MV-HA-VC

The VC fragment corresponding to the split Venus GFP was amplified by PCR with the primers indicated below. The HA epitope was incorporated in the N-terminus within the forward primer. Primers also contained suitable restriction sites to facilitate the subcloning. The PCR product was first subcloned as a PmeI-XhoI fragment in an Actin SV40-polyA attB plasmid of the laboratory. Finally, the HA-VC fragment with the SV40 polyA was subcloned in the MV vector using the PmeI-SpeI restriction sites. Sequence of the plasmid will be provided upon request until the vector is deposited in a public repository.

Forward primer HA-VC: *5’ttaggcggtttaaacgcggccgcgccaccgacgtcatgtacccatacgatgttccagattacgctggggccgcgg ccggggacaagcagaagaacg 3’*

*Reverse primer* VC*: 5’attatagagctcgaggtaccctactattacttgtacagctcgtccatgccgagagtgatccc 3’*

#### MV-Dronc-V5-VC

The Dronc-V5-VC fragment was synthetised by Twist Bioscience. A cDNA wildtype of Dronc was fused to the V5 and VC peptides at the C-terminus. The constructs was subcloned in the MV vector using PmeI-KpnI restriction sites. Sequence of the plasmid will be provided upon request until the vector is deposited in a public repository.

#### MV-wg-RNAi

The construct was built using the primers indicated below. The primers included the inverted repeats for targeting the gene *wingless* previously described in the *Drosophila* Trip collection (HMS00844) (https://fgr.hms.harvard.edu/fly-in-vivo-rnai). The primers were synthetised and subsequently annealed using standard protocols. Then the resulting double strand DNA was subcloned in the MV vector previously opened with Pmel-Kpnl restriction sites.

Forward primer:

*5’ aaaccagttagctcgatatgaatataatatagttatattcaagcatatattatattcatatcgagctagcggtac 3’*

*Reverse primer:*

*5’ cgctagctcgatatgaatataatatatgcttgaatataactatattatattcatatcgagctaactggttt 3’*

### Fly Husbandry and full description of genotypes

All fly strains used are described at www.flybase.bio.indiana.edu unless otherwise indicated. Primary *Drosophila* strains and experiments were routinely maintained on Oxford fly food at 25 °C.

### Full Genotype Description

**Fig. 1**

1b: *Sal-Gal4/MV-HA-VC*

1c: *MV-HA-VC/+; dpp-lexA/+*

1d: *MV-HA-VC/+; Hh-QF/+*

1e: *Sal-Gal4/ MV-HA-VC; Hh-QF/+*

1f: *Sal-Gal4/MV-HA-VC; dpp-lexA/+*

**Fig. 2**

2a: *hs-Flp^122^*; *Sal-Gal4/MV-Dronc-V5-VC*

2b: MV-*Dronc-V5-VC/+; dpp-lexA/hs-mFlp5*

2c: MV-*Dronc-V5-VC /hs-Cre; Hh-QF/+*

**Supplementary Figure 1.**

1b: *Sal-Gal4/ MV-HA-VC; Hh-QF/+*

1d: *Sal-Gal4/ MV-Dronc-V5-VC; dpp-lexA/+*

1e: *Sal-Gal4/ MV-Dronc-V5-VC; Hh-QF/+*

### Immunohistochemistry

Third instar larvae were dissected on ice-cold PBS. The larvae were then fixed and immunostained following standard protocols (fixing in PBS 4% paraformaldehyde, washing in PBT 0.3% (0.3% Triton X-100 in PBS)). Primary antibodies used in our experiments were: anti-GFP (goat, 1:400; Abcam, ab6673); anti-Ptc (1:200, Hybridoma Bank, Apa1); anti-HA (rabbit 1:500, Cell Signalling 3724). The following secondary antibodies were diluted in 0.3% PBT from (Molecular Probes) and used to detect the primary antibodies: anti-goat Alexa 488, anti-mouse Alexa 555 and anti-rabbit Alexa 647. The secondary antibodies were always used at the standard concentration 1:200. DAPI was added to the solution with the secondary antibodies for labelling the DNA in the nuclei (1:1000; Thermo Scientific 62248). After incubation for 2h with the secondary antibodies, samples were washed in PBT several times and mounted in Vectashield.

### Immunoprecipitation and Western Blot

Twenty larvae of each respective genotype were snap-frozen at −80°C, thawed, macerated on ice in 300 μl of RIPA buffer (complemented with protease inhibitor). Then the sample was cleared at 20.000 *g* for 20 min and supernatant toped-up with PBS + 0.3% Triton X-100 to 500 μl. Next the sample was incubated with 15 μl of V5 magnetic beads (chromotek) prewashed three times in PBS + 0.3% Triton X-100 headovenhead at 4°C on a spinning wheel for 90 min. Beads were washed three times in PBS + 0.3% Triton X-100. Bound material was eluted from the beads in 40 μl Laemmli buffer and loaded on a 4–12% gradient gel. Dronc-V5-VC was detected after completing the Western blotting with goat anti GFP antibody (Abcam, 1:2500).

### Imaging of wing discs

Confocal imaging of wing imaginal discs was performed using the Olympus Fluoview FV1200 and the associated software. 55 focal planes were taken per wing disc using the 40x lens. Acquired images were processed using automated Fiji/ImageJ. Generally, Z-stacks were projected, and the channels split. The Reslice function was used to obtain transversal sections of the images. Figures were produced using Adobe photoshop CC2018.

### Generation of genetic mosaics

Larvae *yw hs-Flp1.22*; *MV-Dronc-V5-VC /Sal-Gal4* were heat shocked for 25 mins at 37°C 24-48h after egg laying and dissected at the end of LIII.

Larvae *w; MV-Dronc-V5-VC/+*; *dpp-LexA/hs-mFlp5* were heat shocked for 60 mins at 37°C 48-72h after egg laying and dissected at the end of LIII. We have noticed that the recombination efficiency using the *hs-mFlp5* is lower than the WT and a longer heat-shock treatment is needed to generate the genetic mosaics.

Larvae MV-*Dronc-V5-VC/hs-Cre*; *Hh-QF* were dissected at the end of LIII without giving any heat-shock since the *hs-Cre* line has demonstrated leaky expression^16^. The larvae were maintained at 25°C until dissection at the end of LIII stage.

## Supporting information

Supplementary information Wendler et al

## CONTRIBUTIONS

L.A.B-L. was responsible for the initial conception of the work and the design of all the plasmids described in the manuscript. The empty MV vector, *MV-HA-VC*, *MV-Dronc-V5-VC* and *Beta2-tubulin-Cre* plasmids were generated by L.A.B-L. The experiments shown in Figure 1 and 2 were performed by L.A.B-L. F.W. generated the *Beta2-tubulin-mFlippase-5* and MV-*wg-RNAi*_plasmids. F.W. also performed the immunoprecipitation experiments and Western Blotting described in the manuscript. The transgenic flies shown in the study were generated by S.P. and Bestgene Inc. The initial *in vivo* validation of the MV vector with plasmids not included was made by C.H., A.G., K.R., I.A., Y.S., M.B., S.L., E.S. and G.A. as part of a summer course in molecular biology and fly genetics for undergraduate students. S.K. and L.A.B-L. were responsible for establishing and coordinating the execution of the aforementioned summer course. S.K., FW and L.A.B-L. wrote the manuscript and prepared all the figures of the manuscript. All co-authors have provided useful criticisms and commented on the manuscript before submission.

## DATA AVAILABILITY

All data are incorporated into the article and its online supplementary material. All of the experimental resources generated in this manuscript will be publicly available through different public repositories upon publication and until then will be freely distributed upon reasonable request to the corresponding author. The full sequence of the MV plasmid will be accessible in the Supplementary Text of the manuscript. The plasmids described in the manuscript will be sent to a public repository (e.g. *Drosophila* Genome Resource Repository; https://dgrc.bio.indiana.edu/Home) upon publication. The new *Drosophila* strains described in the manuscript will be submitted to the Bloomington *Drosophila* Stock Center (https://bdsc.indiana.edu/index.html) upon publication.

## ACKNOWLEDGEMENTS

We are grateful to the following investigators for providing flies and reagents: Drs. I. Salecker (for kindly providing all the reagents and flies used in the study related with *mFlippase-5*, and hs-Cre line on the second chromosome), K. Basler (for providing the *Beta2*-tubulin-Flippase WT and *dpp*-lexA strains), Bloomington Stock Center (fly strains), Kyoto Stock Center (fly strains) and DGRC (wild-type cDNAs). We thank Dr. G. Struhl for sharing information regarding the *Btub2* promoter, and Genewiz and Bestgene for making the DNA synthesis and generating the transgenic flies, respectively. We are also grateful for support from the Stanford Bing Overseas Program at Oxford, directed by Dr. S. Sodywola. We also thank the caspaselab members (https://www.caspaselab.com) at Oxford for critical reading of the manuscript and valuable suggestions. This work has been supported by Cancer Research UK C49979/A17516 and the John Fell Fund from the University of Oxford 162/001. L.A.B-L. is a CRUK Career Development Fellow (C49979/A17516) and an Oriel College Hayward Fellow. C.H. is a PhD student on the BBSRC funded Oxford Interdisciplinary Bioscience Doctoral Training Programme (BB/M011224/1). A.G. and F.W are postdoctoral fellows of CRUK (C49979/A17516). K.R.C. was supported by a Vice Provost Undergraduate Education (VPUE) fellowship from Stanford University. Work here was supported by a VPUE Course Development Grant, and by NIH awards (R01 DK107507; R01 DK108817; U01 DK123743 to S.K.K.), the JDRF Northern California Center of Excellence, and by NIH grant P30 DK116074 (S.K.K.) for the Stanford Diabetes Research Center.

