## Supplementary information Wendler et al for "A toolkit to generate inducible and interconvertible *Drosophila* transgenes"

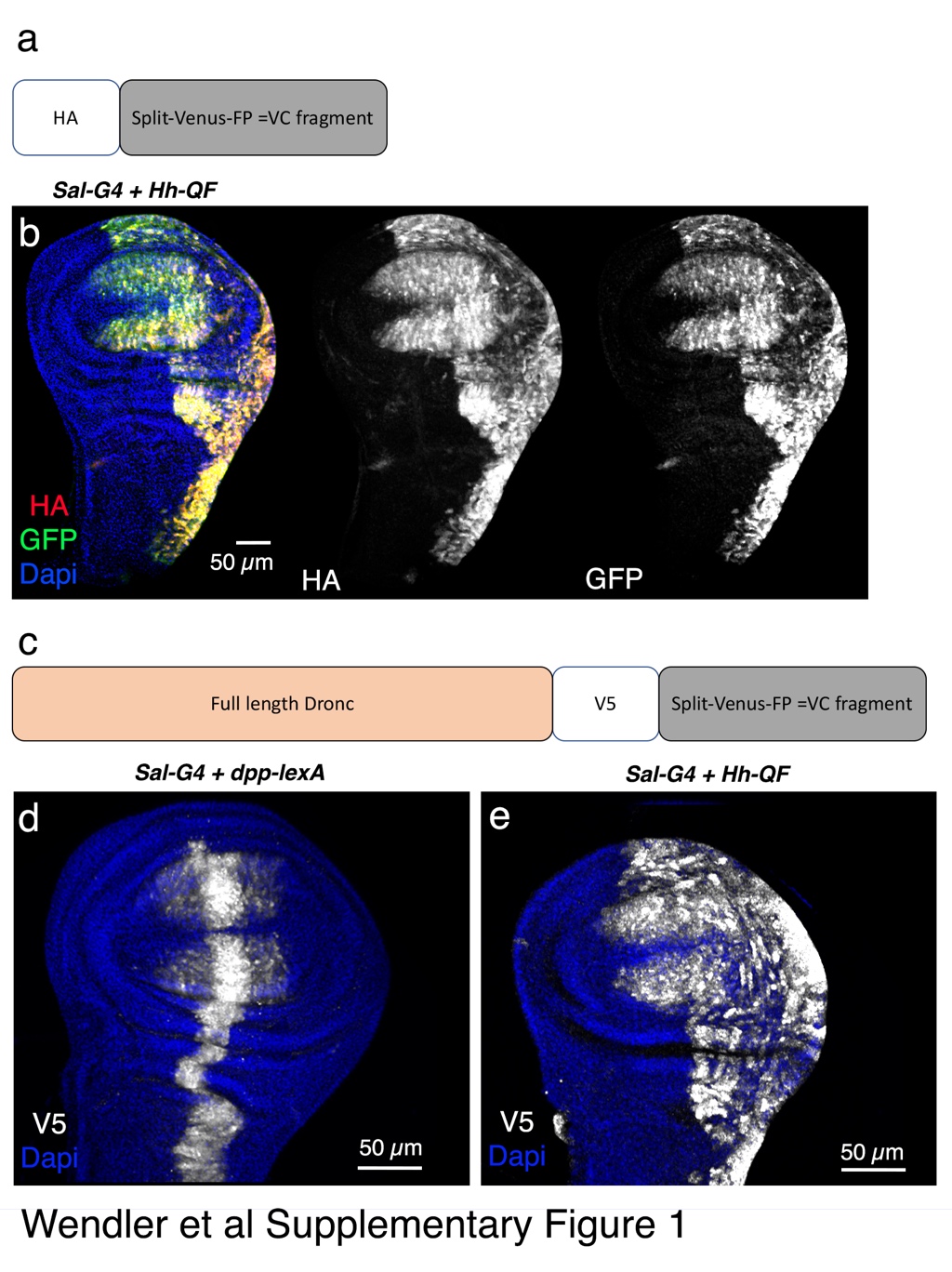

**Supplementary Figure 1**

**a)** Schematic showing the design of the *HA-VC* construct. **b**) Concomitant expression of the HA-VC protein in the *Sal-Gal4* and *Hh-QF* cellular domains of the wing disc; the HA-VC expression was detected by using anti-HA (red) and anti-GFP (green). **c)** Schematic showing the design of the *Dronc-V5-VC* construct. **d-e)** Concomitant expression of the Dronc-V5-VC (grey anti-GFP) in several territories of the wing disc induced by different transcriptional activators. Dapi (blue) stains DNA in the entire figure. Scale bars represent 50 μm in the entire figure.
